# An intestinal cell atlas and organoid model for the threespine stickleback

**DOI:** 10.64898/2026.06.25.734627

**Authors:** Arshad A. Padhiar, Abdelmounaim Nouri, Stefan Keller, Emily Reinhardt, Kathryn Milligan-McClellan, Rebecca Carrier, Natalie C. Steinel, Daniel I. Bolnick, Maria L. Rodgers

## Abstract

The intestine plays a crucial role in physiology, nutrition, and immune function, but intestinal anatomy and cell types have yet to be fully characterized in many fish species, the most diverse group of vertebrates. To address this gap, we characterized the structure and composition of the intestine of threespine stickleback (*Gasterosteus aculeatus*), an emerging model teleost in biological research. Using histology, myeloperoxidase staining, single-cell RNA sequencing, and RNA *in situ* hybridization, we defined major intestinal epithelial, immune, stromal, and stem/progenitor populations. Goblet cells were abundant in proximal and hindgut, while myeloperoxidase-positive granulocytes were evenly distributed throughout the intestine. To facilitate future experimental studies of stickleback intestinal function, we also developed the first intestinal organoid culture from stickleback and show that these cultures recapitulate epithelial architecture and retain expression of canonical intestinal epithelial markers. This organoid platform enables future functional studies of mucosal immunity, host-microbe interactions, and intestinal physiology in stickleback and related teleosts. Together, our integrated approach provides a comprehensive cell atlas and a novel experimental model for studying digestive and immune functions in threespine stickleback.

## Introduction

Vertebrate digestive systems exhibit remarkable diversity in morphology, physiology, and cellular composition, shaped by factors such as diet, microbiome, and pathogen exposure (Karasov and Douglas 2013). This variation is particularly pronounced among fishes, which display a wide range of gut lengths, morphologies, and functional specializations (Wilson and Castro 2010). Some lineages exhibit striking departures from the canonical vertebrate digestive plan, including cyprinid fishes, which lack a stomach entirely (Stroband et al. 1979). Understanding the molecular and cellular bases of this diversity is essential for elucidating the evolutionary forces that shape digestive system function across vertebrates. However, for most fish species, detailed characterization of intestinal cell types, their spatial organization, and functional roles remains limited. This knowledge gap constrains opportunities for comparative and mechanistic studies of digestive physiology and mucosal immunity across the diversity of teleosts, particularly how intestinal cell types contribute to host defense and host–microbe interactions.

Threespine stickleback (*Gasterosteus aculeatus*) have emerged as a powerful model organism for a wide range of biological disciplines, including ecology, evolutionary biology, neurobiology and behavior, genetics, development, toxicology, and immunity. Stickleback have undergone repeated divergence between anadromous marine and freshwater populations following the Pleistocene glacial retreat (McKinnon and Rundle 2002; Reid et al. 2021).

Within freshwater environments, stickleback have adapted to a wide diversity of habitats (e.g., rivers, streams, large oligotrophic lakes, small eutrophic ponds). These habitat transitions are associated with shifts in diet (Stuart et al. 2017; Matthews et al. 2010), gut microbiome composition (Smith et al. 2015), and pathogen exposure (Bolnick et al. 2020), including infection by parasites such as *Schistocephalus solidus* in freshwater populations (Barber and Scharsack 2010). As a result, stickleback are increasingly used to study the proximate mechanisms underlying adaptation, including developmental, immunological, neurobiological, and physiological processes (Norton and Gutiérrez 2019; Strickland et al. 2024), as well as host–microbiome coevolution (Small et al. 2023). In addition, stickleback have emerged as a model for studying peritoneal fibrosis, an immune-mediated response to parasitic infection that can limit parasite growth, with differences in both the extent and resolution of fibrotic responses, providing a potential framework for understanding mechanisms of fibrosis regulation (Hund et al. 2022). Despite this growing interest in stickleback biology, much of the fine-scale anatomy and physiology of the stickleback intestine remains poorly described, limiting the utility of this system for studies of gut function, nutritional homeostasis, and mucosal immunity.

A foundational step toward understanding intestinal function in any species is identifying which cell types are present and how they are distributed along the length of the gut. Key questions include whether cellular composition differs between anterior and posterior regions, and how immune and epithelial cell populations are spatially organized to support mucosal immune function. Traditional histological approaches provide valuable insight into tissue architecture and broad cell classes, but generally lack the resolution needed to distinguish closely related cell subsets, such as specific immune cell types, based on molecular identity. In contrast, single-cell RNA sequencing (scRNAseq) enables the unbiased identification of cell populations based on gene expression profiles, offering detailed insight into cellular composition and potential function within complex tissues (Molla Desta and Birhanu 2025), but scRNAseq typically lacks spatial information. When combined with spatial techniques such as RNA *in situ* hybridization (RNA-ISH), merging these approaches allow transcriptomically defined cell types to be localized within intact tissue (Williams et al. 2022).

Here, we integrate histological analyses, scRNAseq, RNA-ISH, and intestinal organoid culture to define the cellular composition and spatial organization of the threespine stickleback intestine while establishing an experimentally accessible *in vitro* model. While descriptive cellular atlases provide an essential framework, experimental systems that enable functional interrogation of intestinal cell types remain limited in most non-mammalian vertebrates. Three-dimensional intestinal organoids provide a complementary approach by capturing key features of epithelial organization in culture and allowing intestinal epithelial behavior to be studied under controlled conditions (Shin and Kim 2022; Sugimoto and Sato 2017). We therefore establish the first intestinal organoid culture system for this species, extending descriptive insights into an experimentally accessible platform. Together, these approaches generate a comprehensive cellular atlas and a functional toolkit that enhance the utility of stickleback in studying digestive physiology, mucosal immunity, and host-environment interactions in a tractable experimental system.

## Results

### Histological description of the stickleback intestine

H&E-stained sections from the proximal, mid, and hind intestine were clinically assessed by a certified veterinary pathologist (Emily Reinhardt, co-author). Across all segments, the intestinal mucosa exhibited a consistent structural organization, including invaginated villi lined by simple columnar epithelium, a vascular lamina propria, a submucosa, inner circular and outer longitudinal layers of the muscularis, and an outer serosa (Figure S1).

The proximal gut epithelium showed basally positioned nuclei, a prominent apical brush border, and abundant goblet cells. The lamina propria contained capillaries within loose connective tissue, along with scattered lymphocytes and fibroblasts. The submucosa was thin (∼10 μm) and vascularized, while the muscularis consisted of inner circular (∼50 μm) and outer longitudinal (∼25 μm) layers, and the serosa was composed of simple squamous epithelium. The midgut displayed a similar epithelial organization, with fewer lymphocytes and a thicker submucosa (∼20 μm) with a highly vascularized lamina propria. In the hindgut, lymphocytes were rare, with comparable lamina propria and submucosa (∼15 μm), although the muscularis was reduced in thickness.

To quantitatively assess regional specialization along the entire length of the intestine, the gut was prepared using a swiss-roll technique, which allows the full intestinal tract to be visualized in a single histological section. Sections were stained with H&E and Alcian blue to quantify villi length and goblet cell density, respectively. Villi length differed significantly (Kruskal-Wallis test (H(3)=24.2, p<0.05, ε^2^= 0.896)) among regions, with the hind intestine exhibiting the shortest villi relative to the proximal and mid intestine (Figure 1A-F). The mean normalized villi length (villi length(um)/body length(mm)) for the proximal, mid, hind intestine, and rectum were 6.57 ± 0.964, 4.01 ± 0.575, 2.57 ± 0.507, and 3.43 ± 0.230, respectively. Goblet cell density (number of goblet cells/absorption area (mm^2^)) also varied significantly (Kruskal-Wallis test (H(3)=11.2, p<0.05, ε^2^= 0.416)) along the intestine, with the highest density observed in the hind intestine compared to the proximal intestine (Figure 1G-L). The mean goblet cell density for the proximal, mid, hind intestine, and rectum were 962.32 ± 154.553, 1142.29 ± 202.348, 1448.21 ± 182.564, and 1183.02 ± 314.419 cells, respectively. Overall, these results highlight distinct regional specialization along the stickleback intestine, with structural and cellular features suggesting functional differences across gut segments.

**Figure 1.**
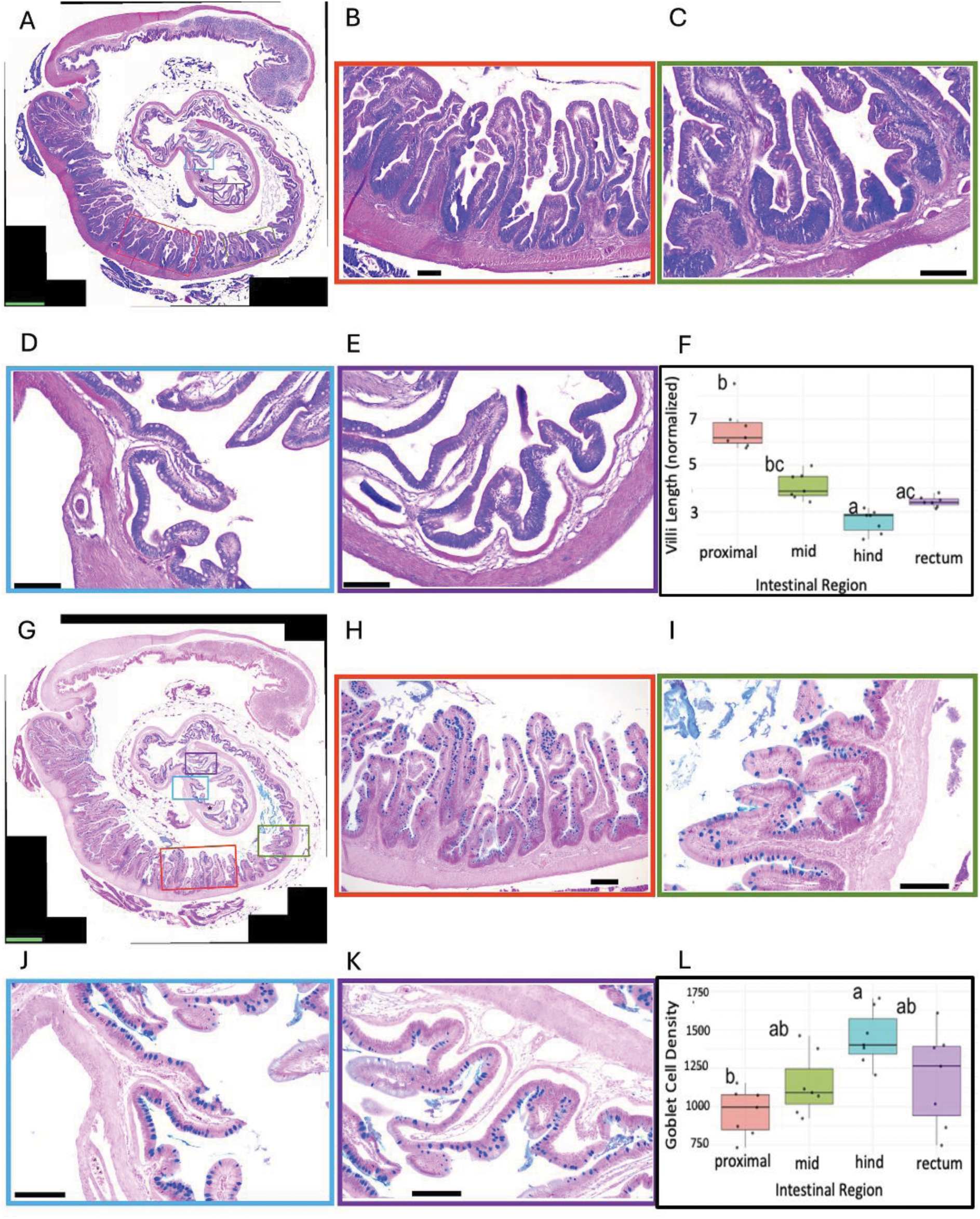
Average villi length and goblet cell density per region along the intestine. (A) Representative stitched image micrograph of the intestinal Swiss roll on H&E stained section. (B-E) Examples of villi from different regions are represented in the colored boxes. (B) proximal intestine, (C) mid intestine, (D) hind intestine (E) rectum. (F) Box and whisker plot of the mean values for each fish per intestinal region, with each point representing the average villi length per individual. (G) Representative stitched image micrograph of the intestinal Swiss roll on Alcian blue stained section. (H-K) Examples of goblet cell distribution along the villi are represented in the colored boxes. (H) proximal intestine, (I) mid intestine, (J) hind intestine (K) rectum. (L) Box-and-whisker plot of the mean values for each fish per intestinal region, with each point representing the average goblet cell density per individual. Significance was calculated using Kruskal-Wallis (P<0.05) followed by Dunn’s post hoc test. Rectal valve (asterisk). Green scale bar = 500um. Black scale bar = 100um.

### Distribution of Myeloperoxidase positive cells along the intestine

To assess the abundance and spatial distribution of innate immune granulocytes in the intestine, cryosections from the proximal gut, midgut, and hindgut were stained using a myeloperoxidase (MPO)-specific histochemical assay. MPO is a peroxidase enzyme abundantly expressed in neutrophils and other granulocytes and is commonly used as a marker for these innate immune cells (Rizo-Téllez et al. 2022). Examining MPO-positive cells therefore provides insight into the distribution of granulocytic immune populations along the intestinal mucosa.

MPO-positive cells were detected throughout all three intestinal regions and were primarily localized within the lamina propria, with occasional cells observed adjacent to the epithelial layer. The overall distribution pattern appeared similar across the proximal gut, midgut, and hindgut (Figure 2A–C). Quantification of MPO staining showed comparable levels across intestinal regions (Figure 2D), with the mean values of 2.38% ± 2.01%, 2.69% ± 2.24%, and 3.44% ± 2.37% in the proximal, mid and hind gut respectively. A one-way ANOVA revealed no significant differences (F_2,24_ = 0.54, P = 0.58), and a linear model treating gut region as an ordered variable showed a non-significant positive trend (β = 0.58, p = 0.28). Together, these findings suggest a largely uniform distribution of MPO-positive granulocytes along the intestine, with only a modest and non-significant increase toward the hindgut.

**Figure 2.**
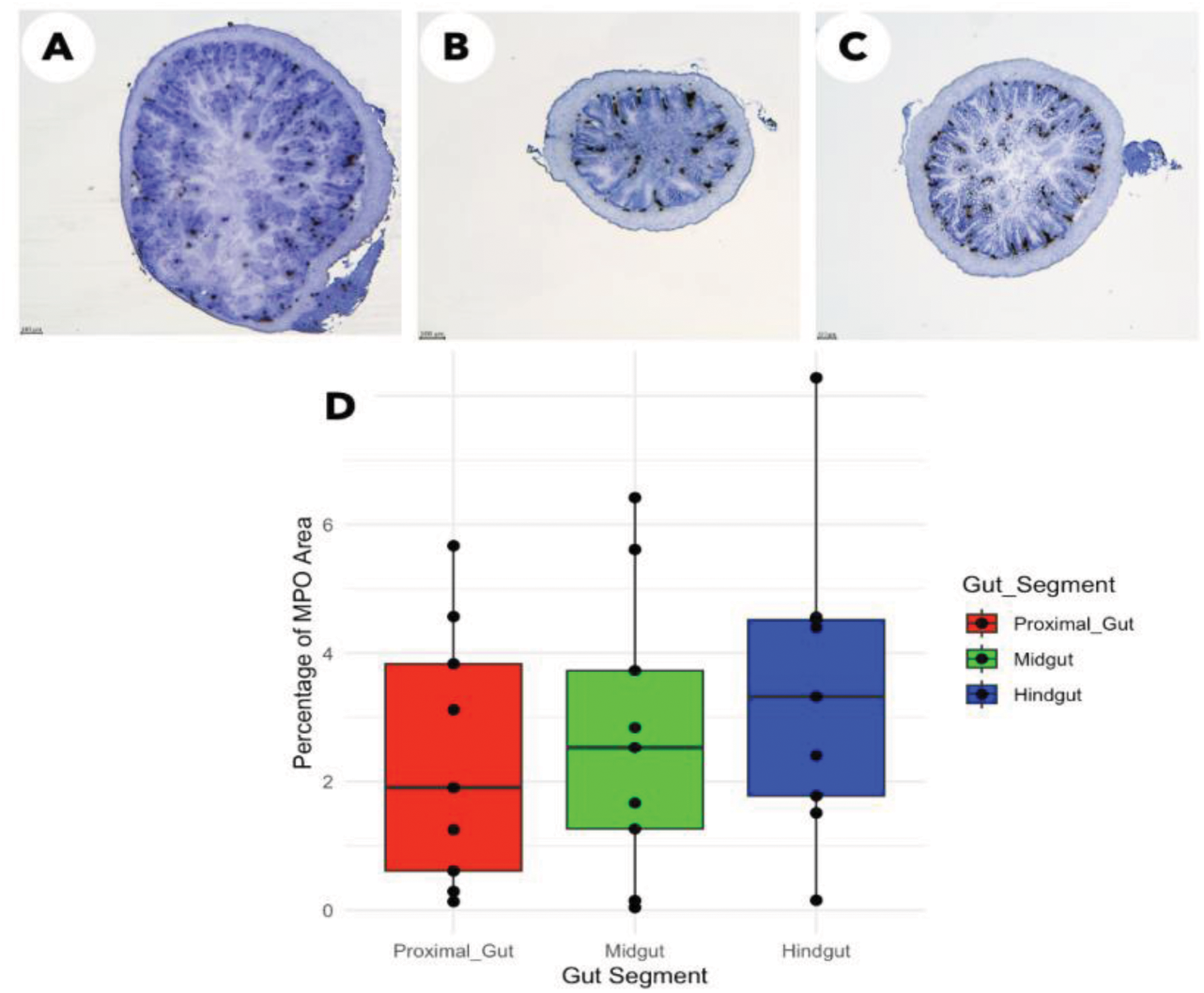
Distribution of myeloperoxidase (MPO)-positive cells along the intestine. (A–C) Representative micrographs showing MPO-positive cells in the proximal gut, midgut, and hindgut, respectively. (D) Box-and-whisker plot showing the abundance of MPO-positive cells across the three intestinal regions. scale bar = 100um

### Single-cell RNA sequencing reveals distinct intestinal cell populations

To characterize the cellular composition of the threespine stickleback intestine at single-cell resolution, we performed single-cell RNA sequencing (scRNA-seq) on dissociated intestinal tissue. Libraries were generated from anterior and posterior intestinal segments processed separately from each of three individuals. We use the resulting data to construct a cell atlas defining the major cell types present in stickleback intestines, and the gene expression patterns that define these. Additionally, we test for differences in cell type composition and gene expression between anterior and posterior halves of the intestine.

Clustering analysis of the combined scRNA-seq dataset identified 23 transcriptionally distinct clusters (Figure 3A and 3B). Twelve of these contained sets of significantly upregulated genes (log2 fold change > 0.5, p < 0.05) and were assigned biological identities based on diagnostic marker gene expression (Table S1). Cluster identities were determined by examining genes significantly upregulated within each cluster and comparing their expression patterns with known markers from teleost and mammalian intestinal lineages, with Ensembl orthology mapping to human and zebrafish via the biomaRt package. The annotated populations included absorptive enterocytes (cluster 20, defined by *anpep*, *mgam*, *tmprss15*, and multiple SLC transporters), goblet cells (cluster 17, defined by *muc5ac*, *spdef*), intestinal stem cells (cluster 6, characterized by rRNA biogenesis and cell-cycle signatures), two distinct cycling/proliferating progenitor populations (clusters 7 and 8) and neuro-immune associated population (cluster 2), (Table S1). In addition, multiple hematopoietic and myeloid populations were identified, including mature erythrocytes (cluster 18, *hbe1, gata2, gfi1b*), proerythroblasts (cluster 19, *lin28b, add2, gmpr*), neutrophils (cluster 21, *mpx, cybb, hvcn1, fpr1*), activated macrophages (cluster 22, *mmp9, il1b, cxcr1*), and a cross-presenting dendritic cell population (cluster 23, *spic, xcr1, il12b*). One cluster (cluster 14) co-expressed markers from multiple distinct lineages including erythroid, pancreatic exocrine, and apolipoprotein gene families; we flag this cluster as a likely doublet or contamination signal. The pancreatic exocrine genes observed in cluster 14 likely reflect contamination from pancreatic tissue.

**Figure 3.**
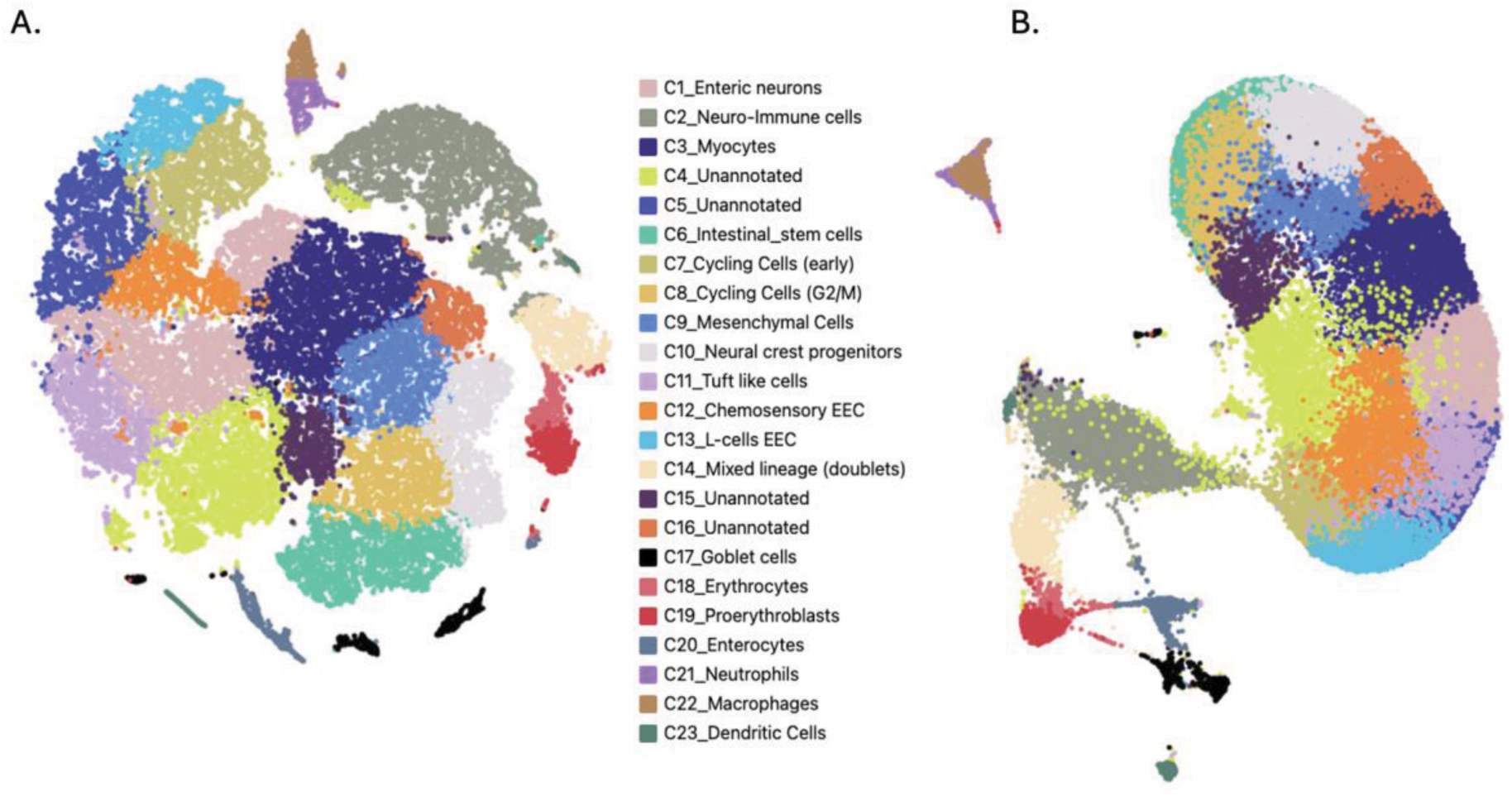
Single-cell RNA sequencing reveals distinct intestinal cell populations. (A) UMAP projection of 41,192 cells from scRNA-seq of the threespine stickleback intestine (N = 3 fish), showing 23 transcriptionally distinct clusters. (B) t-SNE projection of the same dataset.

Unlike mammals, stickleback and other teleosts lack a discrete pancreas; instead, exocrine pancreatic tissue is distributed diffusely along the proximal intestine and associated digestive tissues (Menke et al., 2011). As a result, pancreatic tissue may not have been completely separated from the intestine during sample preparation.

A further seven clusters contained too few cells to yield statistically significant markers individually, but their top genes by fold change aligned with canonical lineage defining signatures and were assigned putative identities. These included enteric neurons (cluster 1, *avp*, *oxt*, *syt15*, *dclk2*), myocytes (cluster 3, *myog*), mesenchymal cells (cluster 9, *col9a1*, *fst*, *myh10*), neural crest progenitors (cluster 10, *hand2*, *fbln7*), tuft-like cells (cluster 11, *dclk1*, *ncam1*, *grm8*), chemosensory enteroendocrine cells (cluster 12, *tas1r2*), and *pyy* positive L-cells (cluster 13, *pyy, htr2b*). The remaining four clusters (4, 5, 15, 16) showed no significant or interpretable differential markers and are reported as unannotated. Together, these annotated and putative populations encompassed the principal epithelial, progenitor, neural/chemosensory, stromal, and immune lineages expected in the teleost intestine (Table S1).

We next compared the single-cell RNA-seq patterns between anterior versus posterior intestines, both at the level of cell type clusters and at the level of single gene transcripts. We observed substantial between-replicate variation in cell type relative abundance (Table S2), and as a consequence no cell cluster exhibited consistently higher abundance in anterior than posterior intestine samples or vice versa (quasibinomial GLM, P > 0.2 for all clusters). A small number of genes were uniquely expressed in either region. To statistically compare expression levels of single genes between intestinal regions, we performed a pseudo-bulk analysis. This merges all the cells’ read counts within a biological sample, to achieve the appropriate level of biological replication (N = 3 fish per region), and is agnostic as to cell type composition and cell clusters. We then used negative binomial GLMs implemented in DESeq2 to find genes with significant differential expression between the three anterior versus posterior intestinal samples (Figure S2). For example, sst1.1(somatostatin 1) was expressed in all three anterior samples and entirely absent in the posterior samples (estimated LFC = 5.33, P_adj_ < 0.00001). Conversely, ENSGACG00000016154 (relaxin-3-like) is more highly expressed in the posterior intestine (LFC = -2.2, P_adj_ < 0.00001), as is fabp6 (LFC = - 3.97, P_adj_ < 0.00001). Relaxin-3 is expressed in enteric neurons in the lamina propria of the mammalian colon. Fabp6 (fatty acid binding protein 6) is a bile transporter regulating gastric secretions in ileal enterocytes of mammals. Hepcidin-like (ENSGACG00000005429) is more abundant in the anterior region (P_adj_ = 0.000019), while ENSGACG00000011851 is more common in the posterior region (P_adj_ = 0.000118). Insulin mRNA (probably derived from pancreatic tissue contamination) was far more abundant in the anterior region (LFC = 6.59, P = 0.00176). Other differentially expressed genes are presented in (Table S3). One especially worth noting is fads2, which is important in fatty acid synthesis. This gene exhibits rapid and parallel evolution of copy number during adaptation to marine versus freshwater, and is marginally more abundant in the anterior intestine (LFC = 0.36, P = 0.00135, though not significant when using adjusted P values). Overall, these analyses did not identify consistent anterior-posterior shifts in cell-type composition, but instead revealed a small set of regionally enriched transcripts.

### Spatial localization of intestinal cell types by RNA *in situ* hybridization

To validate and spatially localize cell populations identified from histology and single-cell transcriptomic analysis, we used multiplex RNAscope on intact stickleback intestinal sections to detect three marker transcripts within the same tissue section. This allowed epithelial and immune-associated populations to be examined in their native tissue context. Positive hybridization signals were detected for multiple markers, confirming the presence and distribution of distinct intestinal cell populations (Figure 4). In one probe set, *neurod1* signal was detected in scattered cells consistent with enteroendocrine identity, while *mpeg1.1* expression marked macrophage-associated cells. *igt* signal was detected at very low frequency and was predominantly observed in a subset of *neurod1*-positive cells, while most *neurod1*-positive cells lacked detectable *igt* signal (Figure 4A). This pattern suggests a low abundance of *igt*-associated (B cells) in the intestinal tissue or may reflect rare co-expression and/or transitional cell states, although signal overlap between channels cannot be excluded.

**Figure 4:**
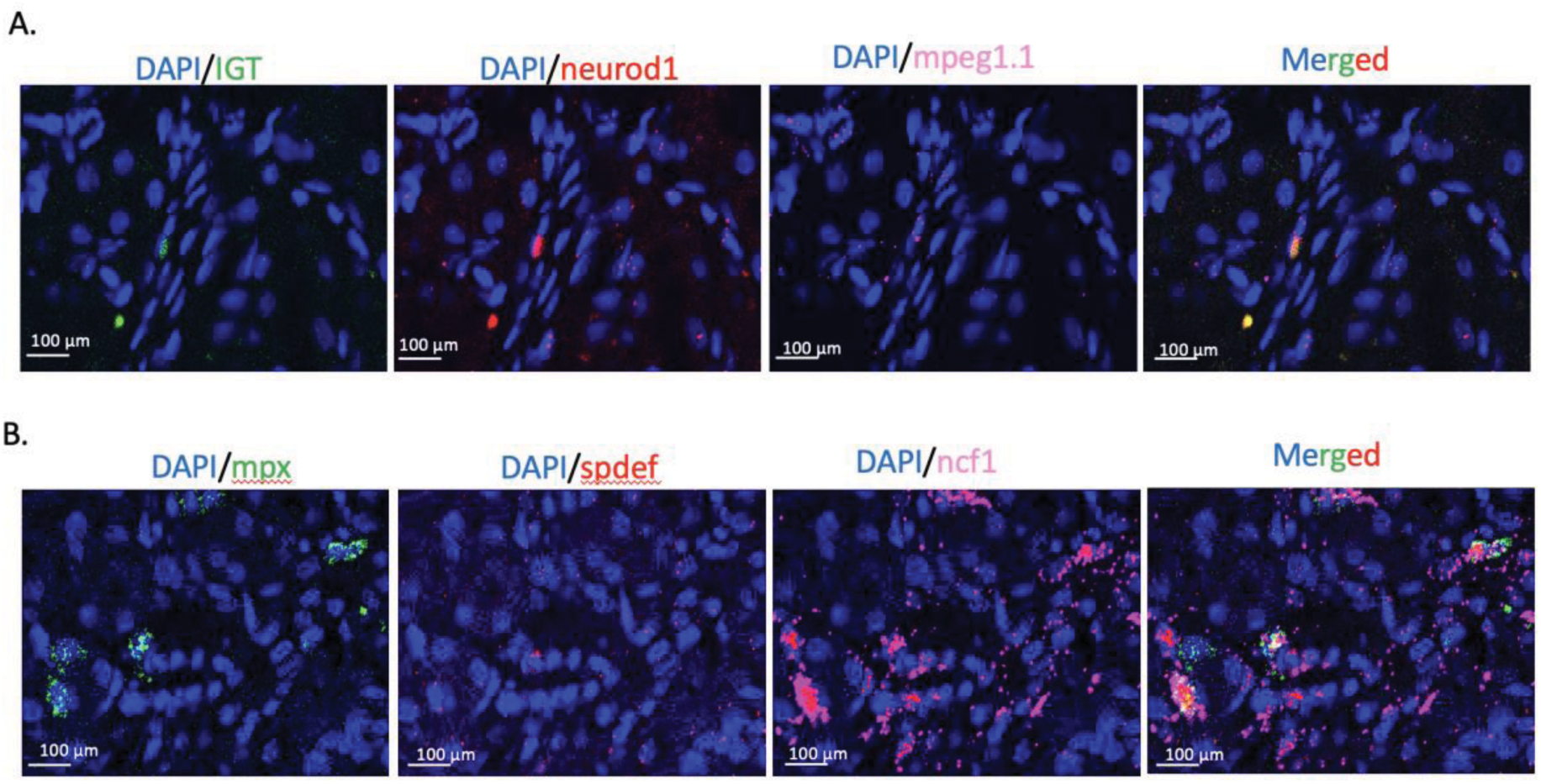
RNA *in situ* hybridization (RNAscope) of stickleback intestinal tissue. RNAscope was used to visualize marker gene expression in intact intestinal sections, with nuclei counterstained by DAPI. (A) *igt* is shown in green, *neurod1* in red, and *mpeg1.1* in pink. *Neurod1* signal was observed in cells consistent with enteroendocrine identity, while *mpeg1.1* expression marked macrophage-associated cells. *igt* signal was limited, with occasional apparent overlap with *neurod1*-positive signal in the merged image; this overlap should be interpreted cautiously because it may reflect close spatial proximity or channel overlap rather than true co-expression. (B) *mpx* is shown in green, *spdef* in red, and *ncf1* in pink. *mpx*-positive signal marked myeloid-associated cells, *spdef* expression marked goblet cell-associated cells, and *ncf1*-positive signal marked neutrophil-associated cells. Signals were primarily localized within the lamina propria.

In a second probe set, *spdef* expression marked goblet cell-associated cells along the intestinal mucosa, while *mpx* and *ncf1* signals identified myeloid/granulocyte and neutrophil-associated populations, respectively (Figure 4B). These immune-associated signals were primarily localized within the lamina propria. Together, these multiplex RNAscope results provide spatial validation of major epithelial and immune-associated cell populations in the stickleback intestine and complement the cellular composition inferred from histology and single-cell transcriptomic analyses.

### Development and transcriptional validation of stickleback intestinal organoids

To extend our characterization of the stickleback intestine into an experimentally tractable *in vitro* system, we established primary intestinal organoid cultures from juvenile threespine stickleback. Intestinal tissue isolated from 30-35 days post-fertilization fish was dissociated and embedded in Matrigel, where epithelial fragments reproducibly formed three-dimensional organoid structures. Primary organoids became visible within 2-3 weeks of plating and developed cystic morphology with a central lumen surrounded by an epithelial layer (Figure 5A). As cultures matured, organoids exhibited budding and crypt-like protrusions, and detached domains were capable of continued expansion, consistent with sustained epithelial growth and proliferative capacity (Figure 5B).

**Figure 5:**
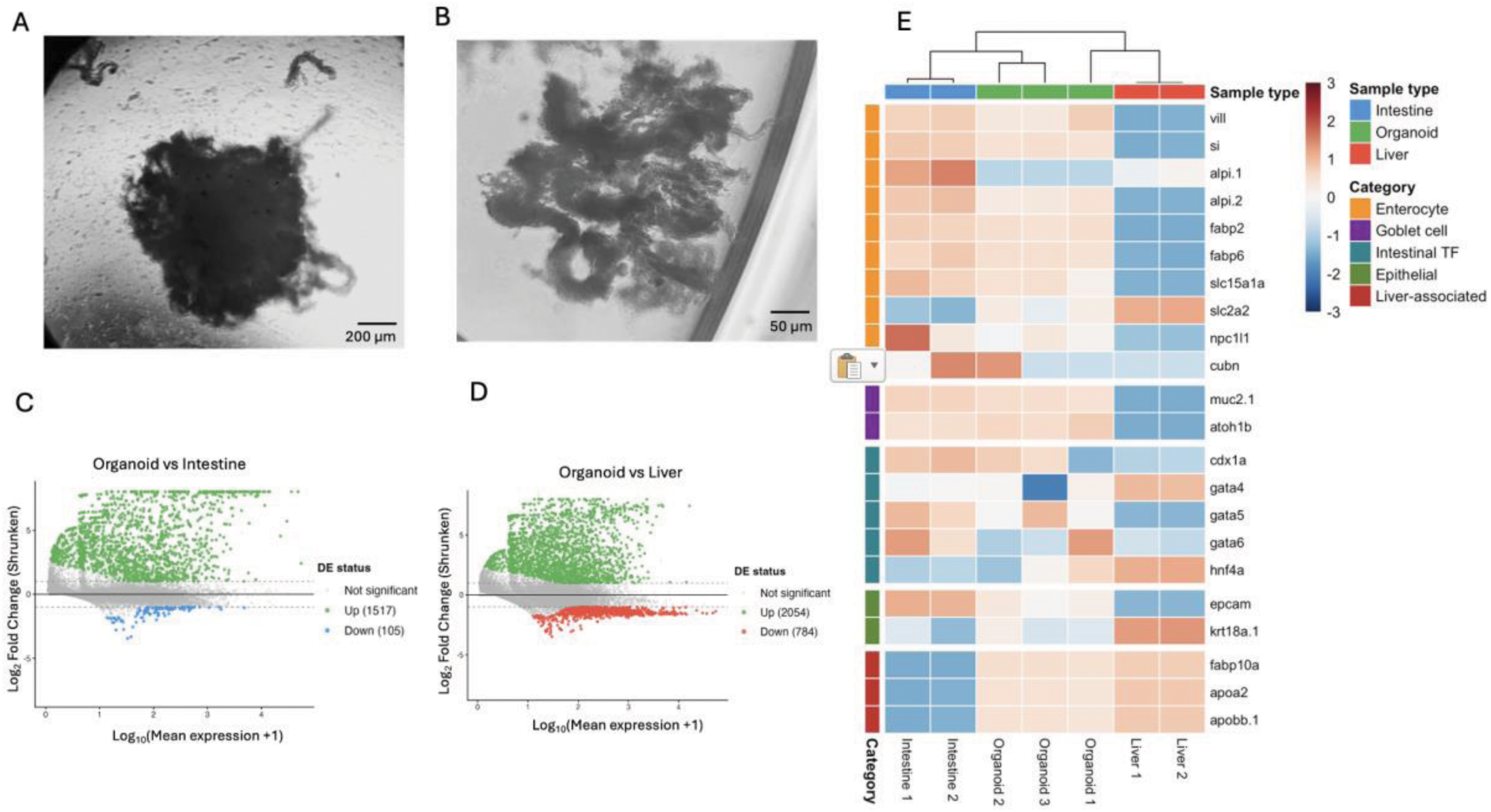
Development and transcriptional characterization of intestinal organoids. (A) Primary intestinal organoids derived from juvenile stickleback, cultured for 5 days. The structures display characteristic 3D cystic morphology with evident crypt-like protrusions (arrowheads), indicative of preserved epithelial polarity and budding activity. (B) Crypt domains have detached from the organoid and initiated autonomous growth outside the Matrigel matrix. The central lumen is surrounded by a continuous epithelial monolayer, from which budding crypt-like domains are emerging, indicating preserved epithelial polarity and proliferative capacity. (C and D) MA plots comparing gene expression between organoids and intestine (left) and organoids and liver (right). Significantly differentially expressed genes are highlighted using adjusted p < 0.05 and absolute shrunken log2 fold change > 1. (E) Heatmap of a canonical intestinal marker panel organized by enterocyte, goblet cell, intestinal transcription factor, epithelial, and liver-associated gene categories. Values are variance-stabilized expression z-scores scaled per gene.

To evaluate whether these cultures retained intestinal molecular identity, we performed bulk RNA sequencing of organoid cultures and compared their transcriptomes with native intestinal tissue and liver, which serves as negative control. Organoid libraries had substantially lower sequencing depth than intestine and liver libraries (Figure S3 A-C), which likely reflected lower RNA yield from organoid cultures. Therefore, downstream interpretation was based on DESeq2-normalized and variance-stabilized expression values.

Despite this limitation, normalized sample-distance analysis showed that organoid replicates remained internally consistent (Supplementary Figure S3-D). Differential expression analysis showed that organoids were transcriptionally distinct from both native intestine and liver, as expected for an *ex vivo* epithelial culture, but showed fewer differences relative to the intestine than to liver (Figure 5C-D, Figure S4). Consistent with this pattern, clustering of organoid-expressed variable genes showed that organoids were more similar to intestine than liver, while maintaining a distinct organoid profile (Supplementary Figure S5). Together, these analyses support a closer transcriptional relationship between organoids and intestinal tissue than between organoids and liver, while also indicating that organoids adopt a distinct culture-associated transcriptional state.

To assess intestinal epithelial identity more directly, we examined canonical intestinal epithelial markers together with liver-associated metabolic genes. Of 24 candidate genes, 22 were detected in the bulk RNA-seq dataset. Organoids expressed multiple intestinal epithelial markers, including enterocyte-associated genes (*fabp2, si, alpi.2, fabp6, slc15a1a, vill*), goblet-associated genes (*muc2.1, atoh1b*), intestinal transcription factors (cdx1a, gata5, gata6), and epithelial markers (*epcam, krt18a.1*) (Figure 5E, Figure S6). Liver-associated metabolic genes (*fabp10a, apoa2, apobb.1*) showed strongest e xpression in liver samples, with lower but detectable expression in organoids. Together, the morphology and bulk RNA-seq data support the intestinal origin of the cultures and validate them as intestinal organoids that retain core intestinal epithelial transcriptional features while adopting a distinct ex vivo expression state.

## Discussion

Fish exhibit extensive diversity in gastrointestinal morphology and physiology, reflecting adaptations to diet, environment, and evolutionary history (Wilson and Castro 2010; Pan et al. 2023; Duque-Correa et al. 2024). In this study, we provide a comprehensive characterization of the threespine stickleback intestine by integrating histological analyses, myeloperoxidase staining, single-cell transcriptomics, RNA *in situ* hybridization, and the first intestinal organoid culture system for this species. Together, these approaches define the cellular composition and spatial organization of the stickleback intestine while establishing new experimental tools for functional studies in this emerging model system. By linking classical histology with molecular and *in vitro* approaches, this work helps bridge descriptive anatomy with mechanistic investigation in a species that has become central to studies of evolution, immunity, and host–microbe interactions.

The intestinal cell types identified in threespine stickleback broadly reflect conserved features of vertebrate gut organization, including diverse epithelial lineages, immune populations, and supporting stromal and muscle-associated cells. The presence of enterocytes, goblet cells, and enteroendocrine cells highlights functional specialization within the epithelium, while immune populations such as macrophages, granulocytes, and monocytes indicate an active mucosal immune compartment. Although fishes display substantial diversity in gut morphology across taxa (Huang et al. 2020), the overall cellular composition observed here is consistent with patterns reported in other teleost species, supporting the idea that core intestinal cell programs are evolutionarily conserved (Lickwar et al. 2017), despite variation in gross anatomy and ecology (Price et al. 2019).

Beyond overall cellular composition, our analyses highlight how intestinal cell populations are organized along the length of the stickleback gut. Histological assessment revealed region-specific differences in villus morphology and goblet cell abundance, while immune cells were distributed throughout the proximal, midgut, and hindgut. The relatively uniform presence of myeloperoxidase-positive granulocytes across gut regions suggests a broadly distributed innate immune surveillance system rather than strong compartmentalization. At the same time, variation in epithelial features such as villus length and goblet cell density points to functional differentiation along the anterior–posterior axis, potentially reflecting differences in nutrient processing, microbial exposure, and local immune demands.

An important strength of this study is the integration of complementary approaches to link cellular identity with tissue context. Single-cell transcriptomic analyses enabled the identification of diverse intestinal cell populations. This scRNA-seq data provided us with cell-type-specific genetic markers. These markers were then used RNA-ISH and histological staining, which provided spatial and morphological context, showing where those cell types are distributed within intact tissue. Rather than relying on a single method, combining these approaches allowed us to cross-validate cell identities and localize major epithelial and immune populations within the intestinal architecture. This integrated framework reduces ambiguity inherent to any one technique and provides a more complete view of intestinal organization in a non-mammalian vertebrate system.

The establishment and transcriptional validation of intestinal organoid cultures from threespine stickleback extends this work beyond descriptive characterization to create an experimentally tractable system. While intestinal organoid models are well developed for mammals, comparable platforms remain limited in fishes, particularly for species with cold-water physiology and distinct epithelial growth requirements (Solhaug et al. 2025).

Optimizing culture conditions for stickleback intestinal epithelium required compatibility with lower temperatures and appropriate medium composition to support epithelial survival and growth. The conditions described here proved to be straightforward, repeatable, and suitable for long-term propagation, cryopreservation, and recovery, providing a practical foundation for future functional studies.

By complementing a cellular atlas with an organoid system, this study creates new opportunities for functional investigation of intestinal biology in stickleback, an emerging model research organism. Organoids provide a controlled platform for probing epithelial responses to microbial signals, immune mediators, and environmental stressors, allowing hypotheses generated from *in vivo* and transcriptomic data to be tested experimentally (Kromann et al. 2024; Wang et al. 2022). In the context of stickleback’s well-characterized ecological divergence and microbiome variation, these cultures offer a means to examine how genetic background and environmental history shape intestinal epithelial function, and interactions with microbial symbionts. More broadly, this framework enables comparative and evolutionary questions to be addressed in ways that are difficult to achieve using whole-animal approaches alone (Yang et al. 2023).

Despite providing a comprehensive framework, this study has several limitations that point to future directions. The cellular atlas was generated under baseline conditions and does not capture how intestinal cell composition or gene expression may shift over development. We focused on intestinal structures in adult fish, and further work should be done across a wider range of ages. We also do not sample multiple populations, having used lab-raised fish descended from a single anadromous population. Further work might profitably compare intestinal architecture across stickleback populations. This is especially relevant because marine and freshwater ecotypes, and populations in different lakes, experience distinct diets, infections, and microbiota (Milligan-Myhre et al. 2016). These ecological differences might drive evolution of distinct intestinal architecture, or induce differences through developmental plasticity. Our analyses were performed in specific-pathogen free, lab-reared animals with comparatively limited microbial communities compared to wild counterparts.

As the microbiome plays an extensive role in intestinal development and physiology, future work should compare intestinal composition, physiology, and gene expression in response to diverse microbial communities. Similarly, while the organoid system establishes a robust epithelial model, it currently lacks immune and microbial components that contribute to intestinal function *in vivo*. Future work integrating additional cell types, co-culture systems, or environmental challenges will further enhance the utility of these tools. These should be replicated across a range of age classes and genotypes. Nonetheless, by combining cellular, spatial, and *in vitro* approaches, this study lays a foundation for mechanistic and comparative studies of intestinal biology in stickleback and other teleost fishes.

## Methods

### Definitions of gastrointestinal segments

We use the terms “proximal gut,” “midgut,” and “hindgut” in this paper, as well as “anterior intestine” and “posterior intestine.” For histological sectioning, the intestine was divided into three equal segments, with the proximal gut adjacent to the stomach, the midgut representing the central portion, and the hindgut connected to the rectum. For single-cell RNA sequencing, the intestine was divided into two equal halves, referred to as the anterior intestine (adjacent to the stomach) and the posterior intestine (adjacent to the rectum).

### Fish Source and Rearing

Laboratory-reared threespine stickleback (Gasterosteus aculeatus) used in this study were derived from parents collected from Sayward Estuary, British Columbia, Canada (50°22′46″N, 125°56′43″W). Wild fish collection was conducted under a scientific permit from the British Columbia Ministry of Lands, Forests, and Environment (Permit #MRVI21-619679), with transportation approved by the Canadian Department of Fisheries and Oceans (License #124591). Fish rearing and euthanasia procedures were approved by the Institutional Animal Care and Use Committees under protocols A21-025 (Bolnick lab) and 21-10-07-Ste (Steinel lab).

### Fish dissection and extraction of the gut

To obtain intestinal tissues for histological analyses, laboratory-reared threespine stickleback (*Gasterosteus aculeatus*) from the Sayward Estuary family were euthanized with an overdose of MS-222 (500 mg/L) and dissected. Euthanized fish were immediately placed on crushed ice. A ventral incision was made from the cloaca to the mandible using sharp surgical scissors, followed by two lateral incisions posterior to the operculum to expose the coelomic cavity. The intestinal tract was excised by making one incision near the stomach and a second incision at the cloaca, then transferred to a Petri dish containing chilled phosphate-buffered saline (PBS).

For cryosection-based regional histology and MPO staining, intestines from nine fish were processed by gently flushing intestinal contents using a gavage needle and syringe. The stomach was removed by cutting at the pyloric caeca junction, and the remaining intestine was divided into three equal segments corresponding to the proximal gut, midgut, and hindgut. Tissue fragments were embedded in water-soluble embedding medium (Tissue-Tek), frozen in a slurry of 2-methylbutane and dry ice for 5 min, and stored at −80 °C until sectioning. Samples were sectioned at 7 µm using a cryostat and mounted onto Superfrost Plus slides. Cryosections were used for regional H&E histology shown in Supplementary Figure S1 and for MPO staining shown in Figure 2.

For whole-intestine swiss-roll histology, a separate set of seven fish was processed before regional subdivision and prepared as described below. Swiss-roll sections were used for H&E and Alcian blue staining shown in Figure 1.

### Swiss-Roll Preparation and Paraffin Embedding

For whole-intestine regional histology, intestines from seven fish were prepared as swiss rolls using a modified protocol based on Kandelouei et al. 2024. Fish were fasted for 24 h before dissection. Briefly, dissected intestines were placed in cold PBS for 30 s and transferred to a Petri dish containing 10% neutral buffered formalin with 5% acetic acid for 30 s. Extraneous visceral fat was removed, and the intestine was transferred to filter paper under a dissecting microscope, straightened, and opened longitudinally from the rectum using fine forceps and microdissection scissors. The lumen was oriented away from the filter paper and handled carefully to minimize damage to the luminal surface. During preparation, tissues were periodically moistened with neutral buffered formalin/acetic acid solution to prevent desiccation and facilitate fixation. Starting from the rectum, intestines were gently rolled lengthwise using fine forceps.

Rolled intestines were transferred to a 6-well plate and initially covered with a small volume of neutral buffered formalin/acetic acid solution for 15 min to allow the tissue to maintain its spiral conformation. Additional fixative was then added, and tissues were fixed at room temperature for 24 h. After fixation, tissues were washed three times in PBS for 5 min, dehydrated through graded ethanol, cleared in xylene substitute, infiltrated with paraffin under vacuum at 55 °C, and embedded in wax molds. Paraffin-embedded samples were stored at 4 °C until sectioning. Tissue samples were sectioned at 7 µm using a microtome, mounted on Superfrost Plus slides, dried overnight at 40 °C, and stored at room temperature until staining.

### Histological Staining of Intestinal Tissue

For regional histological assessment of cryosections, sections from nine fish were thawed at room temperature and fixed in 10% formaldehyde for 20 min. Sections were hydrated through successive ethanol dilutions and stained with Gill’s hematoxylin I (Epredia, 72404, Kalamazoo, MI, USA) for 60 s. Slides were rinsed thoroughly with deionized water, stained with eosin Y solution (Epredia, 6766007) for 30 s, dehydrated through an ethanol series, cleared in xylene substitute, and mounted with coverslips using mounting medium (Epredia, 1900231). These cryosections were used for the regional H&E histology shown in Supplementary Figure S1.

For swiss-roll histology, paraffin-embedded sections from seven fish were deparaffinized in xylene substitute and rehydrated through graded ethanol. Sections were stained with hematoxylin and eosin to assess whole-intestine morphology and villus length, or with Alcian blue to quantify goblet cell density along the intestine. Five sections per fish, distributed along the swiss-roll length, were used for H&E and Alcian blue imaging and quantification.

### Histological Imaging and Quantification

H&E- and Alcian blue-stained swiss-roll sections were imaged using a Leica DM6B microscope, and stitched whole-section images were generated using LAS X software. Images were analyzed in FIJI for regional segmentation and villus length measurements. The rectum was defined as the region distal to the rectal valve. The remaining intestinal length, extending from the rectal valve to the pyloric caeca, was divided into three equal regions corresponding to the proximal intestine, mid intestine, and hind intestine.

Villus length was measured from the base of the epithelium to the tip of the intestinal fold and normalized to body length (n=7, with 5 sections per fish distributed along the Swiss roll length). For branched or complex villi, the longest branch was used for measurement. Goblet cell density was quantified from Alcian blue-stained swiss-roll sections using QuPath (n=7, with 5 sections per fish distributed along the Swiss roll length). Regions of interest were drawn around the villi and lamina propria while excluding the muscular layers, and Alcian blue-positive cells were quantified relative to the analyzed tissue area.

### Myeloperoxidase staining

To assess the abundance and distribution of myeloperoxidase (MPO)-positive granulocytes across the intestine, cryosections from the proximal gut, midgut, and hindgut were obtained from nine threespine sticklebacks (*Gasterosteus aculeatus*) from the Sayward Estuary family. Five spaced cryosections were prepared from each gut segment and processed according to the manufacturer’s instructions using an MPO staining kit (Sigma-Aldrich, 390A).

The sections were first fixed at ambient temperature for 30 seconds in a formaldehyde–ethanol solution to preserve tissue morphology. After fixation, the slides were rinsed with distilled water and allowed to desiccate in the dark. The slides were then immersed in a freshly prepared reaction buffer containing peroxidase indicator in 1× Trizmal buffer supplemented with 3% hydrogen peroxide. The sections were incubated for 20 min in the dark at 20 °C. Following incubation, the slides were rinsed under gently flowing tap water and allowed to dry.

Subsequently, the slides were counterstained with hematoxylin solution for one minute to visualize cell nuclei. The stained sections were then coverslipped and stored in an opaque box at room temperature until microscope imaging. Imaging was performed using a Leica DM6B upright microscope equipped with a DMC5400 color CMOS camera. Image analysis was conducted using FIJI software, and MPO-positive staining was quantified as the percentage of stained surface area relative to the total surface area of each gut section.

### Single-cell RNA sequencing

Single-cell libraries were generated from the intestines of three individual laboratory-reared threespine stickleback (*Gasterosteus aculeatus*) from the Sayward Estuary family described above. Anterior and posterior intestinal segments were processed separately from each individual for single-cell RNA sequencing.

### Generation of single-cell suspensions

To generate single-cell suspensions, stickleback were euthanized with MS-222 and the intestines were removed. Each intestine was divided into two equal portions representing the anterior and posterior halves. Each tissue piece was placed on ice in a sterile 24-well plate containing 2 mL of R-90 medium (90% RPMI 1640 with L-glutamine and without phenol red, supplemented with 10% distilled water).

Each intestinal segment was cut longitudinally and further minced into small fragments. Tissue dissociation was performed mechanically using a sterile pipette tip. The resulting suspension from each well was passed individually through a 40 µm nylon mesh filter (BD Falcon), and an additional 2 mL of chilled R-90 medium was added. Cell suspensions were centrifuged at 550 × g for 10 min at 4 °C. The supernatant was discarded, and the cell pellet was resuspended in 2 mL of chilled R-90 medium. This wash step was repeated once, after which cells were resuspended in 1 mL of chilled R-90 medium. Cell suspensions were transported on ice to The Jackson Laboratory for Genomic Medicine (Farmington, CT) for library preparation and sequencing.

### Single-cell library preparation and sequencing

Within one hour of dissection, cells were washed and resuspended in PBS containing 0.04% BSA. Cell viability was assessed using a Countess II automated cell counter (Thermo Fisher Scientific). Approximately 12,000 cells were loaded onto a single lane of a 10x Genomics Chromium Controller. Single-cell capture, barcoding, and library preparation were performed using the 10x Genomics 3′ Gene Expression kit (v3.1 chemistry), following the manufacturer’s protocol (CG000204) (Zheng et al. 2017).

cDNA and final libraries were assessed for quality using an Agilent 4200 TapeStation and quantified by KAPA qPCR. Libraries were sequenced on an Illumina NovaSeq 6000 platform, targeting approximately 6,000 recovered cells per library with an average sequencing depth of 50,000 read pairs per cell.

### scRNA-seq data processing and analysis

Illumina base call files were converted to FASTQ format using bcl2fastq v2.20.0.422. Reads were aligned to a reference genome constructed from the 2020v5 *G. aculeatus* assembly and annotation files available at https://stickleback.genetics.uga.edu/ (Nath et al. 2020). Ensembl annotations (release 95) were combined with repeat, Y chromosome, and revised annotations from Nath et al. using AGAT v0.4.0 (Dainat et al. 2021). A STAR-compatible reference genome was generated using Cell Ranger v6.0.0 (10x Genomics).

The Cell Ranger count pipeline was used to generate gene-by-cell count matrices for each library. Downstream analyses were performed using Scanpy v1.3.7 and the Loupe Cell Browser (Wolf et al. 2018). Cells with fewer than 400 detected genes, more than 20% mitochondrial RNA content, or more than 500 hemoglobin transcripts were excluded from further analysis. Filtered count matrices were concatenated, normalized by total counts per cell, and log-transformed.

Highly variable genes (n = 3,500), identified based on dispersion (Zheng et al. 2017; Satija et al. 2015), and were used for principal component analysis. Batch effects were corrected using Harmony (Korsunsky et al. 2019). Batch-corrected principal components were used to construct a neighborhood graph with 40 nearest neighbors, followed by dimensionality reduction using UMAP (McInnes et al. 2018). Clustering was performed using the Leiden algorithm (Traag et al. 2019), with additional subclustering applied as needed to resolve visually distinct populations. Final embeddings and clustering metadata were imported into the Loupe Cell Browser using Cell Ranger aggr v3.1.0 for interactive visualization. Libraries contained between 12,278 and 19,984 cells, with mean reads per cell ranging from 13,173 to 35,512 and median detected genes per cell ranging from 257 to 779.

### Marker analysis and cell-type annotation

Differential gene expression between each cluster and all remaining cells combined (one-vs-rest) was computed in the Loupe Browser and exported as a complete gene table (17,740 genes; all detected genes were retained, with no pre-filtering applied during export). Marker analysis was performed in R v4.6.0. Significant cluster markers were defined as genes with one-vs-rest log2 fold change > 0.5 and unadjusted p-value < 0.05. Twelve of the 23 clusters met these thresholds and were classified as annotated. For clusters with fewer than ten significant markers under these strict thresholds, likely reflecting reduced statistical power due to small cluster sizes, the top 30 genes ranked by positive log2 fold change (irrespective of p-value) were inspected for canonical lineage-defining markers. Seven such clusters were assigned putative cell-type labels based on convergent canonical signatures (e.g., cluster 3 as myocytes based on MYOG; cluster 11 as tuft-like cells based on DCLK1 and NCAM1; cluster 13 as L-cells based on PYY). The remaining four clusters showed no significant differential markers in either direction and were retained but left unannotated.

Stickleback gene identifiers were cross-referenced against the Ensembl annotation database for Gasterosteus aculeatus (assembly GAculeatus_UGA_version5) using the biomaRt R package (Durinck et al. 2009). For each gene, the following attributes were retrieved: official gene symbol, description, gene biotype, chromosome, human ortholog with orthology confidence type, and zebrafish ortholog with orthology confidence type. Annotations were returned for 12,818 of 17,740 input gene identifiers (72.3%). The remaining identifiers were retained in the analysis but lacked orthologue data and Ensembl annotation in the current assembly. Orthology confidence type was used to flag ambiguous calls (one2many, many2many) during manual review of cluster marker lists. Cell-type assignments were made through manual curation of the top-ranked markers per cluster and cross-referenced against published intestinal cell-type markers from teleost and mammalian literature.

### Anterior versus posterior comparison

Single-cell RNA sequencing data were obtained from both anterior and posterior intestinal samples derived from the same three individuals. We used quasibinomial models to test, for each cell cluster, whether anterior versus posterior intestinal samples differed in the relative abundance of each cell cluster. We used the Seurat package in R (Satija et al. 2015) to implement pseudobulk pooling of cells within each biological replicate sample, then used DESeq2 to implement a general linear model testing whether each gene is consistently more expressed within one gut region.

#### RNAscope

To understand where specific cell types were localizing along the intestinal tract, we chose six gene targets to examine expression patterns related to cell types: *igt* (B cells), *neurod1* (enteroendocrine cells), *mpeg1.1* (macrophages), *mpx* (myeloid cells), *spdef* (goblet cells), *ncf1* (neutrophils). Probes for each of these genes were designed by Bio-Techne/Advanced Cell Diagnostics. We used formalin-fixed paraffin embedded tissues (procedure outlined in “Fish dissection and extraction of the gut” followed by thawing and immersing sections in a 10% formaldehyde solution for 20 minutes to fix the tissue) and used the RNAscope HiPlex12 Reagents Kit (488, 550, 650) v2 according to the manufacturer’s instructions.

Imaging was performed with a Nikon A1R confocal scope at the University of Connecticut’s Advanced Light Microscopy Facility.

### Generation and culture of intestinal organoids

Intestinal organoids were generated from juvenile stickleback (30–35 days post-fertilization) as described in a public protocol by (Padhiar et al. 2026). Briefly, fish were euthanized in 0.5 g/L MS-222 (pH 7.4) and the intestinal tract was dissected under a stereomicroscope on a 2 % agar plate. Gut contents were removed by gentle squeezing, and tissues from approximately 10 individuals were pooled. Mucus was cleared with 0.1 M DTT in PBS containing 3× antibiotic-antimycotic, followed by enzymatic digestion in 1.4 Wünsch units/mL Liberase TM and 24 U/mL DNase I for 45–60 min at room temperature. Digested tissue was filtered through a 70 μm strainer, centrifuged, and the crypt-enriched pellet was resuspended in Corning Matrigel. Matrigel domes (35 μL per well of a 24-well plate) were overlaid with IntestiCult Organoid Growth Medium (Mouse) (STEMCELL Technologies) supplemented with 1× antibiotic-antimycotic. Cultures were maintained at 25 °C without CO₂, with medium changes every 2–3 days. Organoids were passaged every 10–14 days using Gentle Cell Dissociation Reagent (STEMCELL Technologies) and reseeded in fresh Matrigel. For long-term storage, organoids were cryopreserved in a 1:1 mixture of CryoStor CS10 and IntestiCult medium and stored in liquid nitrogen.

## Acknowledgements

We gratefully acknowledge the contribution of JAX-UConn Single Cell Genomics Center, comprising the Single Cell Biology service, Genome Technologies and Cyberinfrastructure high performance computing resources at The Jackson Laboratory, for expert assistance with the work described herein. These shared services are supported in part by the JAX Cancer Center (P30 CA034196). We specifically acknowledge William Flynn and Martine Seignon for bioinformatic assistance. We also gratefully acknowledge the contribution of the Advanced Light Microscopy Facility at the University of Connecticut for their expert assistance with the work described herein, particularly Christopher O’Connell. This work was supported by a Gordon and Betty Moore Foundation Aquatic Symbiosis grant to DIB and NCS and RC (GBMF9323), and an NIH R01 to DIB (1R01AI123659-01A1), and a NIH R01 to NCS (R01AI146168)

## Author Contributions

Conceptualization of project: MLR, AAP, DIB, AN, NCS, and KM-M

Formal Analysis: AAP, MLR, AN, ER, SK, and MS

Funding Acquisition: DIB, NCS, and KM-M

Investigation: AAP, MLR, SK, and AN

Writing – Original Draft Preparation AAP, MLR, SK, and AN

Writing – Review & Editing

AAP, MLR, AN, DIB, NCS, SK, ER, KM-M, and MS

## Supplementary Materials

The Supplementary Materials file contains Supplementary Figures S1-S6 and Supplementary Tables S2-S3. Supplementary Table S1 is provided separately as an Excel workbook containing scRNA-seq cluster annotations.

