## Supplementary material for "An intestinal cell atlas and organoid model for the threespine stickleback": Figure S1-S6, and Table S2-S3

**Supplementary Figures**

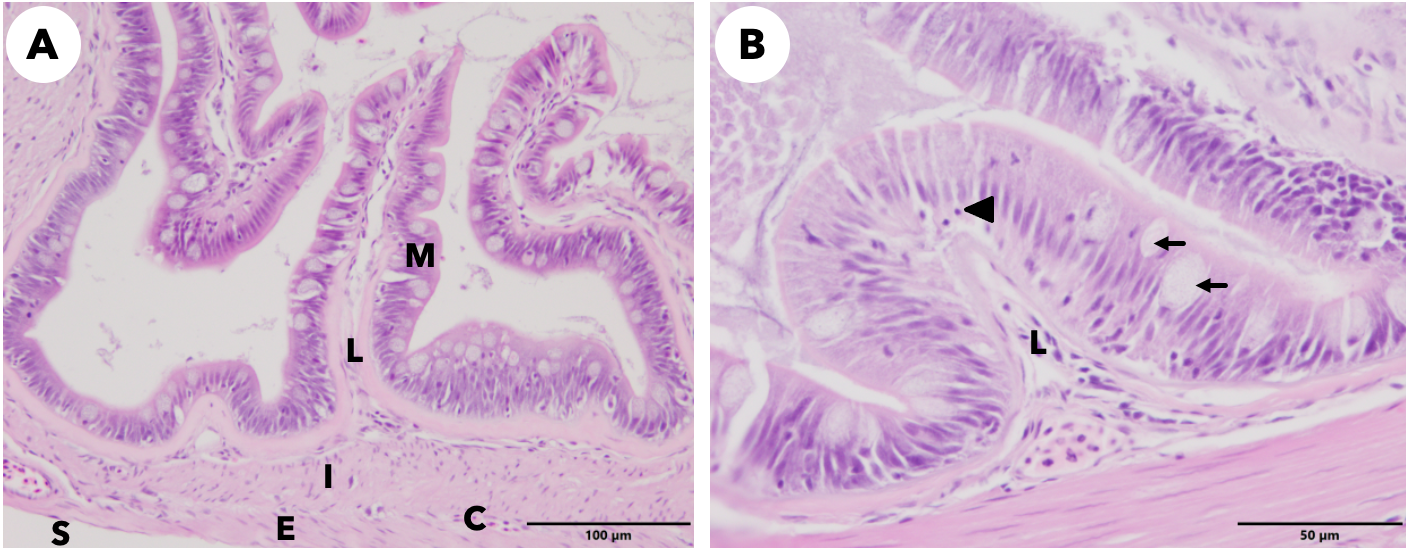

**Figure S1. Histological organization of the stickleback intestine (H&E).**
(A) Cross-section of the gut showing the mucosa (M), lamina propria (L), inner muscular layer (I), outer muscular layer (E), serosa (S), and a capillary (C).
(B) Higher-magnification view of the mucosa (M) showing goblet cells (arrows) and a lymphocyte (arrowhead).

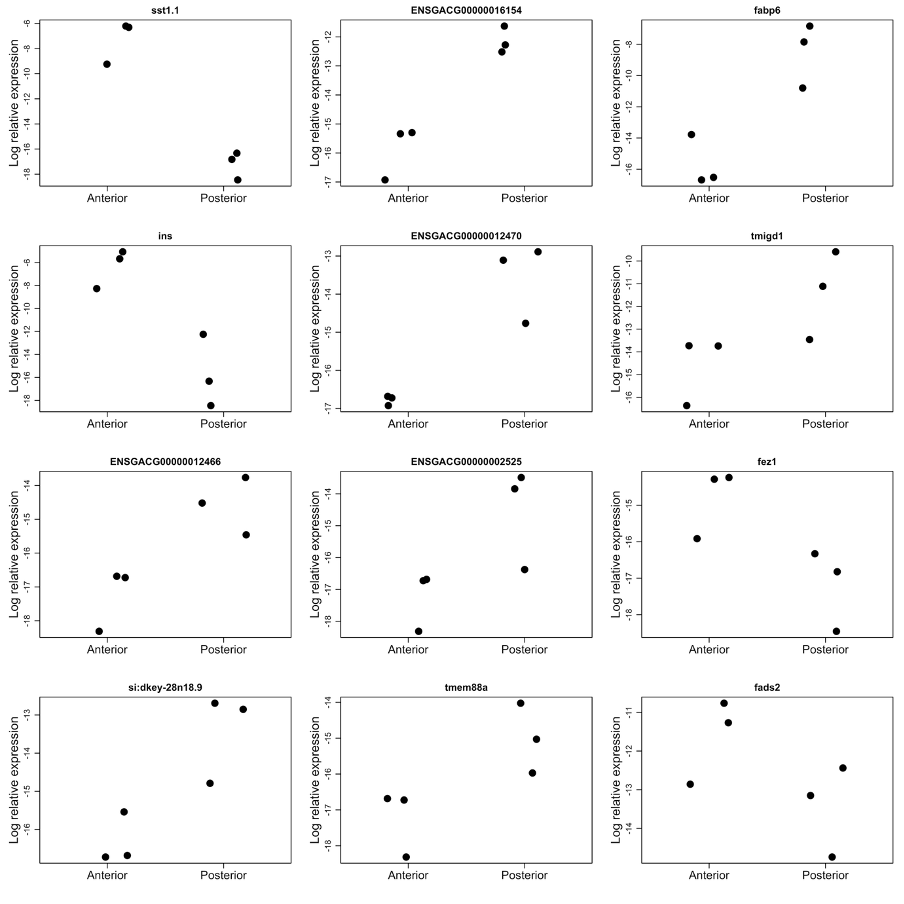

**Figure S2. Regionally enriched transcripts in anterior and posterior intestine.**

Expression levels of selected genes differentially expressed between anterior and posterior intestinal samples based on pseudo-bulk DESeq2 analysis. Each point represents one biological replicate (N = 3 per region).

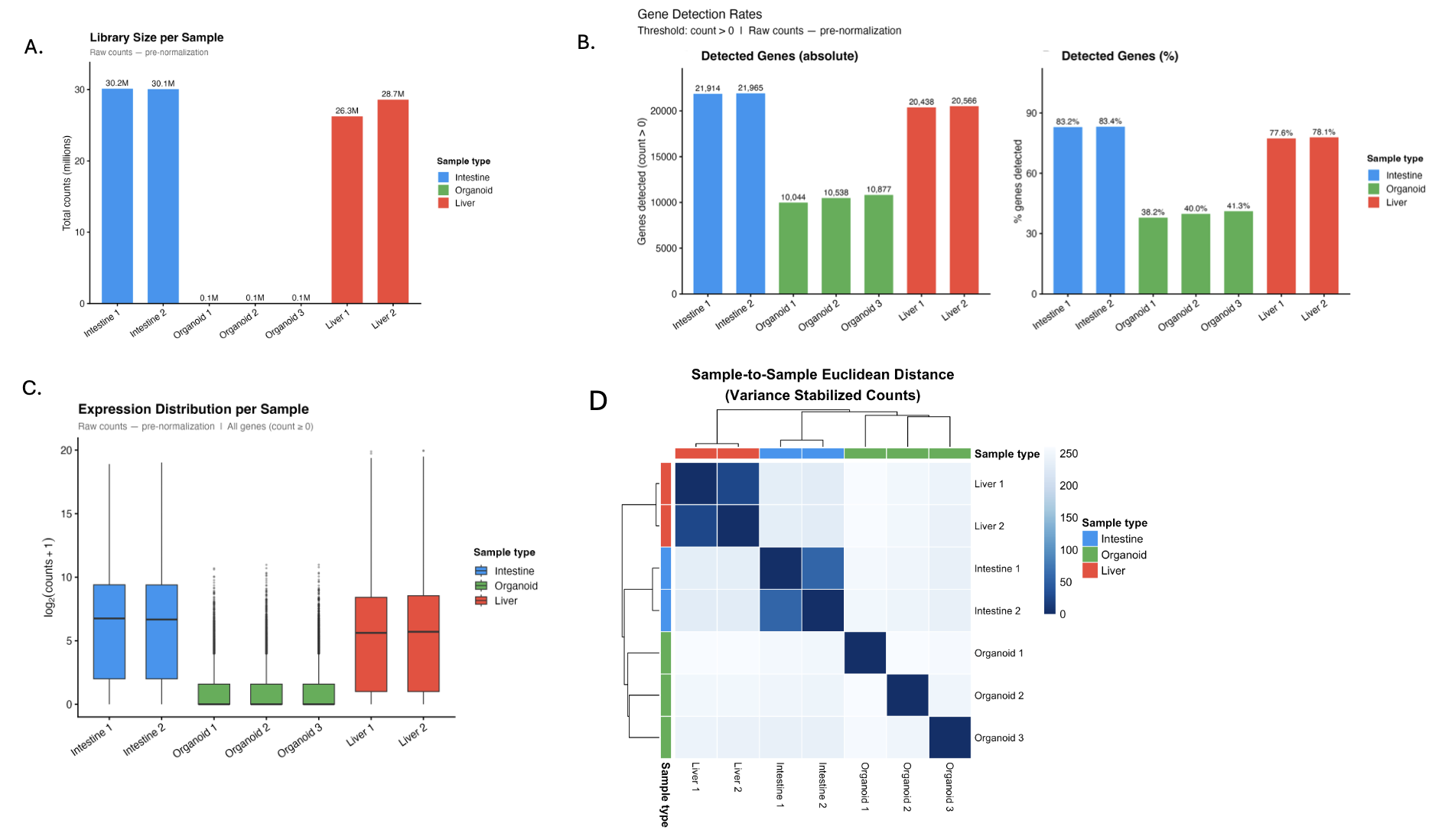

**Figure S3. RNA-seq quality control and normalized sample relationships for organoid, intestine, and liver libraries.**

(A) Total raw counts per sample before normalization. Organoid libraries had substantially lower sequencing depth than intestine and liver libraries. (B) Gene detection rates shown as both the absolute number of genes detected and the percentage of genes detected per sample using a count > 0 threshold. (C) Raw-count expression distributions shown as log2(counts + 1) across all genes. (D) Sample-to-sample Euclidean distance heatmap calculated from variance-stabilized counts. Lower distances indicate greater similarity between samples. Together, these analyses show reduced sequencing depth and gene detection in organoid libraries, while normalized sample relationships still show reproducible clustering of organoid samples.

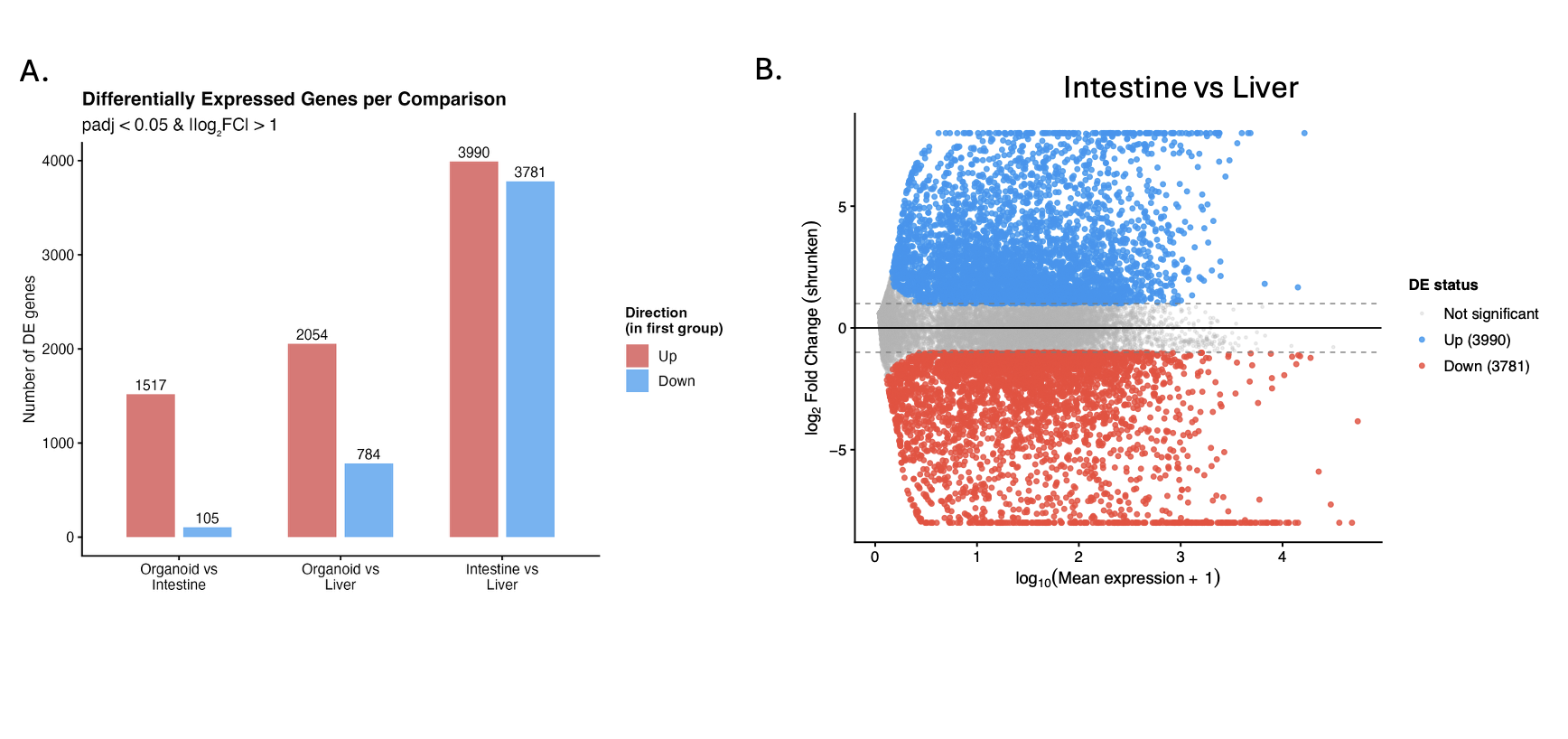

**Figure S4. Differential expression summary.**
(A) Number of differentially expressed genes for each pairwise comparison using adjusted p < 0.05 and absolute shrunken log2 fold change > 1. Up and down indicate genes with higher or lower expression in the first group named in each comparison. (B) MA plot comparing native intestine and liver as a tissue-level reference comparison.

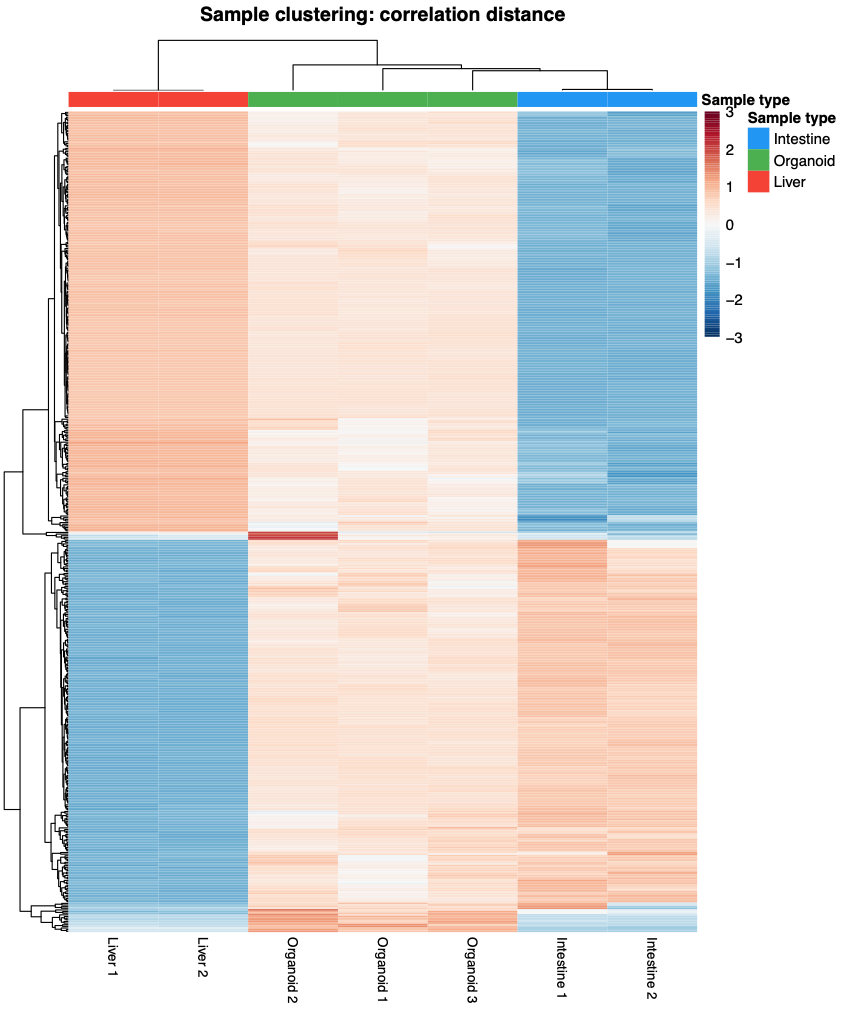

**Figure S5. Organoid-expressed global expression heatmap.**
Heatmap of the top 500 most variable genes among genes expressed in organoids, defined as mean organoid VST expression > 7. Rows show z-score scaled expression values per gene, and columns show samples clustered by correlation distance. This exploratory analysis shows that liver samples remain distinct from the intestine-organoid expression program while organoids retain a distinct culture-associated profile.

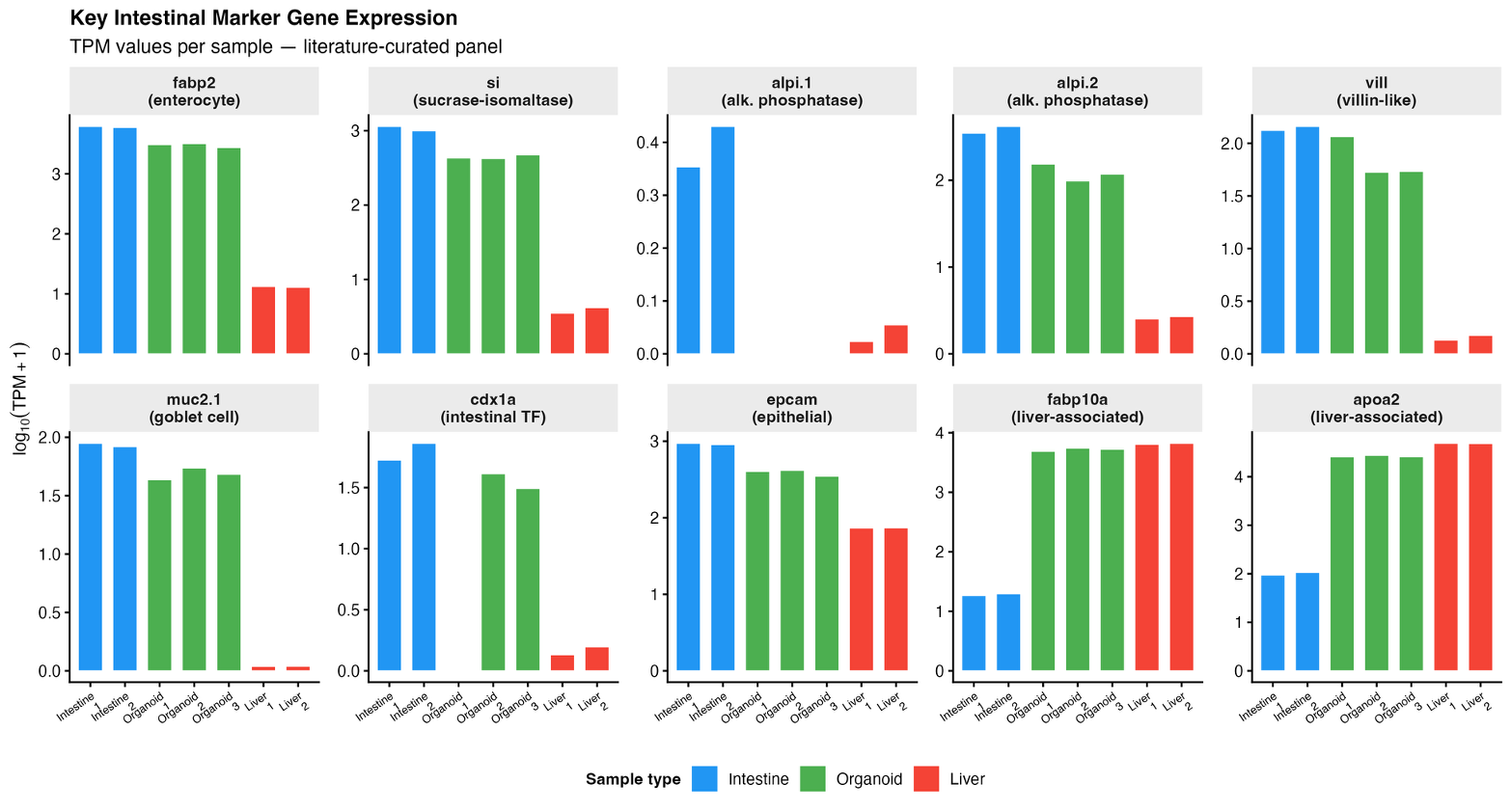

**Figure S6. Representative marker gene expression.**
Bar plots showing log10(TPM + 1) expression for selected intestinal epithelial and liver-associated genes across intestine, organoid, and liver samples. These plots provide gene-level support for the marker heatmap shown in the main figure.

**Supplementary Table**

| Cell cluster | Anterior 1 | Anterior 2 | Anterior 3 | Posterior 1 | Posterior 3 | Posterior 4 | Average % cells |
| --- | --- | --- | --- | --- | --- | --- | --- |
| 1 | 61 | 118 | 26960 | 46 | 7292 | 86 | 21.3 |
| 2 | 251 | 9590 | 307 | 8956 | 1 | 248 | 25.9 |
| 3 | 6560 | 3 | 868 | 6 | 22 | 7014 | 15.0 |
| 4 | 7 | 2 | 92 | 8 | 10252 | 4 | 8.9 |
| 5 | 336 | 105 | 13 | 22 | 7 | 9383 | 8.9 |
| 6 | 125 | 1921 | 725 | 654 | 451 | 148 | 4.5 |
| 7 | 3528 | 8 | 0 | 12 | 3 | 86 | 4.6 |
| 8 | 303 | 603 | 219 | 175 | 116 | 621 | 2.2 |
| 9 | 832 | 82 | 170 | 416 | 94 | 383 | 2.3 |
| 10 | 0 | 0 | 877 | 0 | 684 | 4 | 1.1 |
| 11 | 404 | 71 | 58 | 677 | 64 | 192 | 1.8 |
| 12 | 303 | 275 | 250 | 197 | 119 | 232 | 1.5 |
| 13 | 189 | 278 | 126 | 93 | 90 | 256 | 1.1 |
| 14 | 135 | 49 | 112 | 137 | 124 | 115 | 0.7 |
| 15 | 74 | 39 | 12 | 45 | 12 | 61 | 0.3 |
| Total | 13108 | 13144 | 30789 | 11444 | 19331 | 18833 |  |

**Supplementary Table S2**. Comparison of anterior versus posterior intestinal cell cluster abundance. Each entry in the table is the number of cells assigned to a given cluster, within a particular sample.

| **Gene** | **P-value** | **Log2 fold change** | **Adjusted P value** |
| --- | --- | --- | --- |
| sst1.1 | 3.034808e-26 | 5.33196924 | 4.872991e-22 |
| ENSGACG00000016154 | 2.586283e-14 | -0.22227202 | 4.152795e-10 |
| fabp6 | 2.861339e-11 | -3.97304564 | 4.594453e-07 |
| gcgb | 2.337692e-10 | 0.48040937 | 3.753632e-06 |
| ENSGACG00000005429 | 1.188568e-09 | 0.05383993 | 1.908484e-05 |
| ENSGACG00000011851 | 7.341870e-09 | -0.04981537 | 1.178884e-04 |
| ins | 1.096954e-07 | 6.59243015 | 1.761379e-03 |
| ENSGACG00000012470 | 2.925050e-06 | -0.06708065 | 4.696752e-02 |
| tmigd1 | 1.454465e-05 | -0.85234187 | 2.335434e-01 |
| ENSGACG00000012466 | 8.531905e-05 | -0.02291003 | 1.000000e+00 |
| rgs5a | 9.929508e-05 | 0.32342782 | 1.000000e+00 |
| ENSGACG00000000423 | 1.381857e-04 | 0.06158893 | 1.000000e+00 |
| ENSGACG00000002525 | 1.726629e-04 | -0.03272825 | 1.000000e+00 |
| ptgs2b | 2.195843e-04 | 0.03237525 | 1.000000e+00 |
| dok4 | 2.637283e-04 | 0.03410428 | 1.000000e+00 |
| fez1 | 3.477049e-04 | 0.01832553 | 1.000000e+00 |
| si:dkey-28n18.9 | 8.192395e-04 | -0.08079418 | 1.000000e+00 |
| pyyb | 1.094876e-03 | 1.38920188 | 1.000000e+00 |
| tmprss15 | 1.098829e-03 | 0.02347963 | 1.000000e+00 |
| fads2 | 1.356033e-03 | 0.36160329 | 1.000000e+00 |

**Supplementary Table S3**. Anterior versus posterior intestine gene expression differences, as estimated from a DESeq model applied to pseudo-bulk data (N = 3 fish per intestinal region).
